# Crystal structures of the sheeppoxvirus encoded inhibitor of apoptosis SPPV14 bound to Hrk and Bax BH3 peptides

**DOI:** 10.1101/2020.03.10.986331

**Authors:** Chathura D. Suraweera, Denis R. Burton, Mark G. Hinds, Marc Kvansakul

**Affiliations:** La Trobe Institute for Molecular Science, Department of Biochemistry and Genetics, La Trobe University, VIC 3086, Australia; Bio21 Molecular Science and Biotechnology Institute, The University of Melbourne, Parkville, Australia

**Keywords:** Poxvirus, sheeppoxvirus, apoptosis, X-ray crystallography, isothermal titration calorimetry, Bcl-2

## Abstract

Programmed death of infected cells is used by multicellular organisms to counter viral infections. Sheeppoxvirus encodes for SPPV14, a potent inhibitor of Bcl-2 mediated apoptosis. We reveal the structural basis of apoptosis inhibition by determining crystal structures of SPPV14 bound to BH3 motifs of proapoptotic Bax and Hrk. The structures show that SPPV14 engages BH3 peptides using the canonical ligand binding groove. Unexpectedly, Arg84 from SPPV14 forms an ionic interaction with the conserved Asp in the BH3 motif in a manner that replaces the canonical ionic interaction seen in almost all host Bcl-2:BH3 motif complexes. These results reveal the flexibility of virus encoded Bcl-2 proteins to mimic key interactions from endogenous host signalling pathways to retain BH3-binding and pro-survival functionality.

## Introduction

The altruistic death of infected host cells via programmed cell death or apoptosis is a potent feature of the first line of defence against invading pathogens [1]. The ability of multicellular hosts to eliminate infected cell via apoptosis forced viruses to evolve sophisticated strategies to counter host cell apoptotic defences, including the use of virus encoded homologs of the B-cell lymphoma 2 or Bcl-2 family [2]. As primary modulators of the mitochondrially mediated or intrinsic apoptosis pathway the Bcl-2 family can be divided into two factions; the prosurvival members and the proapoptotic members [3]. All mammalian Bcl-2 family members are characterized by the presence of one or more of the Bcl-2 homology or BH sequence motifs. The prosurvival members, which include Bcl-2, Bcl-w, Bcl-x_L_, A1, Mcl-1 and Bcl-b all feature multiple BH motifs as well as a transmembrane region that enables localization to the outer mitochondrial membrane [4]. The proapoptotic family members can be further separated into two groups, the multi BH motif proteins Bak, Bax and Bok, and the BH3-only proteins Bim, Bid, Puma, Bad, Bik, Bmf, Hrk and Noxa that only feature the BH3 motif [5]. Mechanistically, the BH3 only proteins act by either relieving the ability of prosurvival Bcl-2 to hold Bak and Bax in an inactive state, or may directly activate Bak and Bax [6]. The interplay between the members is mediated by a helix-in-groove interaction, where a helical BH3 motif is bound in the canonical hydrophobic ligand binding groove on prosurvival Bcl-2 proteins [7]. Ultimately, it is the balance of the prosurvival and proapoptotic Bcl-2 members that determines the fate of a given cell.

Large DNA viruses encode a number of sequence and structural homologs of Bcl-2, including examples among the *adenoviridae* [8], *asfarviridae* [9, 10], *herpesviridae* [11, 12] and *iridoviridae* [13, 14]. Among the *poxviridae*, the majority of genera have been shown to harbor one or more examples, including the *orthopoxviridae* [15, 16], *leporipoxviridae* [17, 18], *cervidpoxviridae* [19, 20], *avipoxviridae* [21, 22], *parapoxviridae* [23] and *chordopoxviridae* [24], with the vaccinia virus F1L and myxomavirus M11L the prototypical members of the family. Amongst the *capripoxviridae*, sheeppoxvirus has been shown to encode the potent apoptosis inhibitory protein SPPV14, a Bcl-2 homolog [25]. SPPV14 has been shown to inhibit apoptosis in transfected cells after treatment with etoposide, arabinose-C or UV irradiation. Furthermore, interaction studies revealed that SPPV14 is able directly bind BH3 motif peptides of several mammalian proapoptotic Bcl-2 family members, and functionally replaces F1L in the context of a recombinant vaccinia virus infection. However, the structural and biochemical basis for apoptosis inhibition by SPPV14 remained unclear. Here we report the crystal structures and affinity data of SPPV14 bound to BH3 motifs of proapoptotic Bcl-2 family members. Our findings provide a structural and biochemical platform to understand sheeppoxvirus mediated inhibition of apoptosis.

## Materials and methods

### Protein expression and purification

Recombinant SPPV14ΔC31, (uniprot accession number A0A2P1A8X8; SPPV14 residues 1-145 with a 31 residue C-terminal deletion and hereafter referred to as SPPV14) was expressed and purified as previously described [25]. SPPV14 mutants R84A and Y46A were synthesized as codon optimized cDNA (Genscript), cloned into the bacterial expression vector pGEX-6P3 and purified as previously described [25]. Purified protein obtained using glutathione affinity chromatography was then subjected to size-exclusion chromatography using a Superdex S75 10/300 or 16/600 column mounted on an ÄKTA Pure system (GE Healthcare) equilibrated in 25 mM HEPES pH 7.5, 150 mM NaCl and 10 mM TCEP (Tris(2-carboxyethyl)phosphine hydrochloride), and fractions analysed using SDS-PAGE. The final sample purity was estimated to be greater than 95% based on SDS–PAGE analysis and appropriate fractions were pooled and concentrated using a centrifugal concentrator with 3 kDa molecular weight cut-off (Amicon^®^ Ultra 15) to final concentration of 10.0 mg/ml. Recombinant SPPV14ΔC31:Bim BH3 complex was subjected to size exclusion chromatography as described above for SPPV14ΔC31 alone. Molecular weight standards (Sigma-Aldrich) used for column calibration were: bovine thyroglobulin (670 kDa), bovine γ-globulin (158 kDa), chicken ovalbumin (44 kDa), horse myoglobin (17 kDa), vitamin B12 (1.350 kDa).

### Measurement of dissociation constants

Binding affinities were measured by isothermal titration calorimetry (ITC) using a MicroCal iTC200 system (GE Healthcare) at 25°C using wild type truncated SPPV14 as well as two mutants SPPV14 Y46A and R84A in 25 mM HEPES pH 7.5, 150 mM NaCl, 10 mM TCEP at a final concentration of 30 μM. BH3 motif peptides were used at a concentration of 300 μM and titrated using 19 injections of 2.0 μl of ligand solution. All affinity measurements were performed in triplicate. Protein concentrations were measured using a Nanodrop UV spectrophotometer (Thermo Scientific) at a wavelength of 280 nm. Peptide concentrations were calculated based on the dry peptide weight after synthesis. The BH3-motif peptides used were commercially synthesized and were purified to a final purity of 95% (GenScript) and based on the human sequences previously described [26] except for Bok BH3: VPGRLAEVCAVLLRLGDELEMIRPSV (accession code Q9UMX3, residues 59-84).

### Crystallization and structure determination

Crystals for SPPV14:Hrk BH3 and SPPV14:Bax BH3 complexes were obtained by mixing SPPV14 with human Hrk 26-mer or Bax 28-mer peptide using a 1:1.25 molar ratio as described [27] and concentrated using a centrifugal concentrator with 3 kDa molecular weight cut-off (Amicon^®^ Ultra 0.5) to 10.0 mg/ml. Concentrated protein was immediately used for crystallization trials. Initial high throughput sparse matrix screening was performed using 96 well sitting drop trays (swissic, Neuheim, Switzerland).

SPPV14:Hrk BH3 crystals were grown by the sitting drop vapour diffusion method at 20°C in 0.2 M sodium potassium tartrate, 0.1 M bis-tris propane pH 6.5, 20% PEG 3350. The crystals were flash cooled at −173 °C in mother liquor supplemented with 25% glucose. The SPPV14: Hrk BH3 complex formed single cuboidal crystals belong to space group C222_1_ with a=43.28 Å, b=70.66 Å, c=115.08 Å, α=90.00°, β=90.00°, γ=90.00° in the orthorhombic crystal system.

All diffraction data were collected at the Australian Synchrotron MX2 [28] beamline using an Eiger detector with an oscillation range 0.1° per frame at a wavelength of 0.9537 Å. The diffraction data were integrated using XDS [29] and scaled using AIMLESS [30]. A molecular replacement solution was obtained using the Balbes [31] automated pipeline and identified M11L (PDB code: 2BJY)[32] as the optimal model. SPPV14:Hrk BH3 crystals contained one molecule of SPPV14 and one Hrk BH3 peptide in the asymmetric unit, with a 43% solvent content and gave final TFZ and LLG values of 32.2 and 1539, respectively. The final model of SPPV14:Hrk BH3 was built manually over several cycles using Coot [33] and refined using PHENIX [34] giving final R_work_/R_free_ values of 0.221/0.235, with 99% of residues in the favoured region of the Ramachandran plot and no outliers.

SPPV14:Bax BH3 crystals were obtained in 1.0 M lithium chloride, 0.1 M citrate pH 5.0, 30% PEG 6000. The crystals were flash cooled at −173 °C in mother liquor. The SPPV14: Bax BH3 complex formed single rod-shaped crystals belong to space group P2_1_ with cell dimensions a=100.27 Å, b=78.56 Å, c=107.76 Å, α=90.00°, β=110.96°, γ=90.00° in the monoclinic crystal system. Diffraction data collection, integration and scaling were performed as described above. The molecular replacement was performed using PHASER [35] with the previously solved structure of SPPV14:Hrk BH3 as a search model. SPPV14:Bax BH3 crystals contain eight molecules of SPPV14 and eight Bax BH3 peptides, with a 48.9% solvent content and final TFZ and LLG values of 11.2 and 1720 respectively. The final model of SPPV14:Bax BH3 was built manually as described above giving final R_work_/R_free_ values of 0.238/ 0.288, with 99% of residues in Ramachandran favoured region and no outliers. All images for SPPV14:Hrk and SPPV14:Bax complexes were generated using PyMOL molecular graphic system version 1.8.6.0 (Schrödinger, LLC, New York, USA). The coordinates for the structures were deposited at the Protein Data Bank (PDB) using accession codes 6XY4 and 6XY6. All raw images were deposited at the SBGridDB using their PDB accession codes [36]. All software were accessed through the SBGrid suite [37].

### Bioinformatics analyses

The closest structural homologs of SPPV14 were identified using the Dali server [38] by searching the entire PDB. Protein interaction interfaces were examined using the ePDB PISA (Proteins, Interactions, Surfaces, Assemblies) program [39]. Sequence alignments were performed using MUSCLE [40].

## Results

We previously showed sheeppoxvirus encoded SPPV14 binds peptides spanning the BH3 motif of the proapoptotic Bcl-2 family members Bak and Bax as well as Bim, Bid, Bmf, Hrk and Puma as determined by surface plasmon resonance (SPR) [25]. To determine the BH3 peptide binding specificity we performed ITC employing a recombinant protein, SPPV14ΔC31 (hereafter referred to as SPPV14), lacking the 31 C-terminal residues of the predicted transmembrane motif and BH3 motif peptides derived from human proapoptotic Bcl-2 proteins (Table 1). ITC confirmed BH3 peptides of Bak and Bax as well as Bim, Bid, Bmf, Hrk and Puma as high affinity interactors, as previously reported [25]. In addition, we observed weaker affinities for BH3 peptides of Bad and Bok, with K_D_ values of 5.2 μM and 7.6 μM, respectively. To understand the structural basis for BH3 motif binding by SPPV14 we expressed and purified recombinant truncated SPPV14 and reconstituted SPPV14:Hrk BH3 and SPPV14:Bax BH3 complexes. Crystals of the SPPV14:Hrk BH3 complex diffracted to 2.05 Å (Table 2). After molecular replacement using M11L as a search model (PDB ID 2BJY) [18] clear and continuous electron density was observed for SPPV14 residues 5-139 and Hrk residues 26-50 (Figure 1A), with the remaining residues presumed disordered. SPPV14 adopts a globular helical bundle structure comprising a central helix α5 around which 6 additional helices are scaffolded to form the classical Bcl-2 fold (Figure 2A). A Dali structural similarity analysis identified deerpoxvirus DPV022 (PDB ID: 4UF3) [20] as the closest structural homolog of SPPV14 with an rmsd of 2.1 Å over 106 Cα atoms, with M11L the second closest match with an rmsd of 2.3 Å over 111 Cα atoms (Figure 2C).

**Figure 1:**
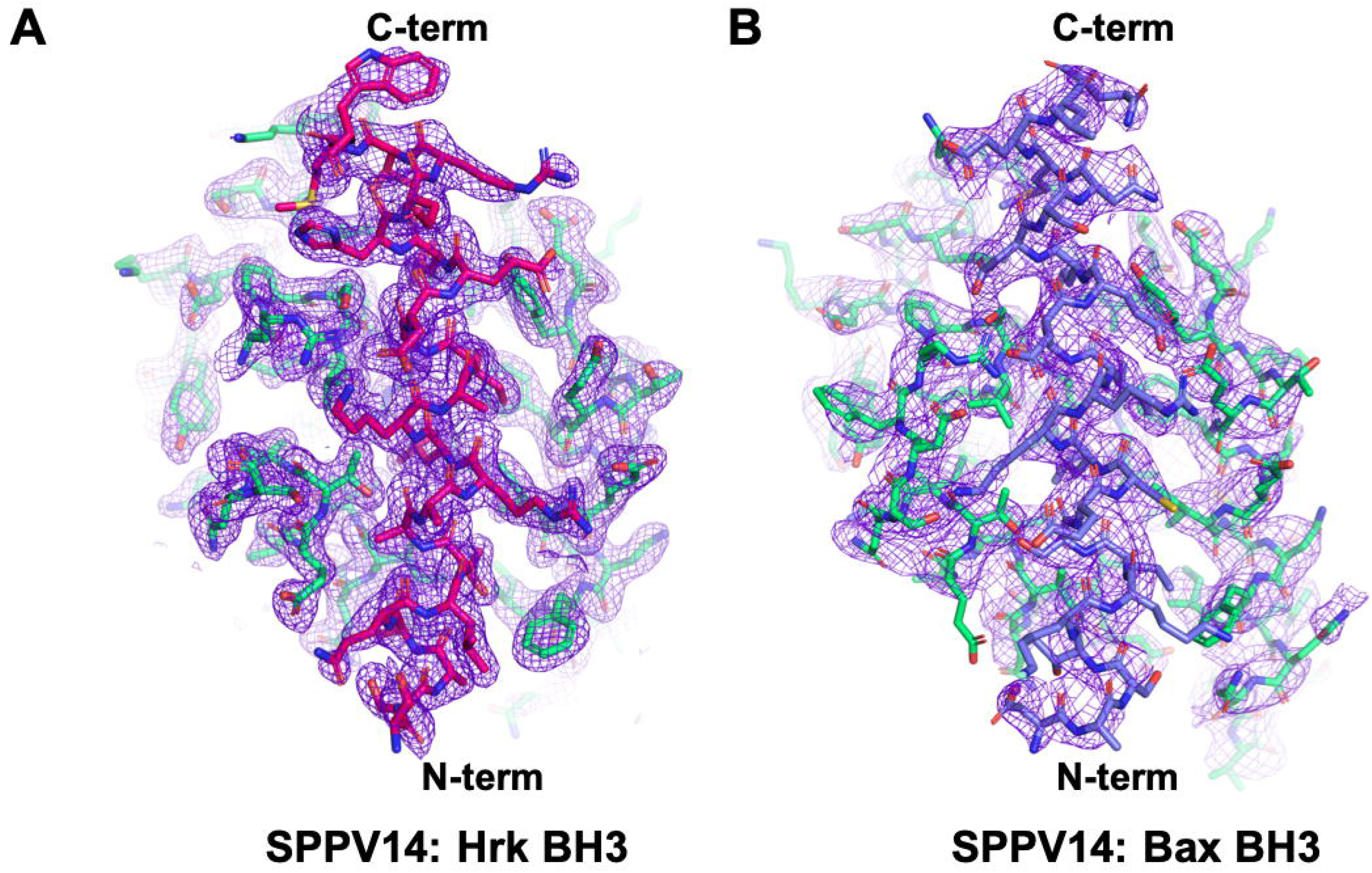
Electron density maps of SPPV14:Hrk and Bax BH3 complexes. **(A)** 2Fo-Fc electron density maps of SPPV14:Hrk BH3 complex interface contoured at 1.5 σ. SPPV14 is shown as green sticks and Hrk as pink coloured sticks. **(B)** 2Fo-Fc electron density maps of SPPV14:Bax BH3 complex interface contoured at 1.5 σ. SPPV14 is shown as green sticks and Bax as slate coloured sticks. The N and C-termini of the BH3 peptide are labelled.

**Figure 2:**
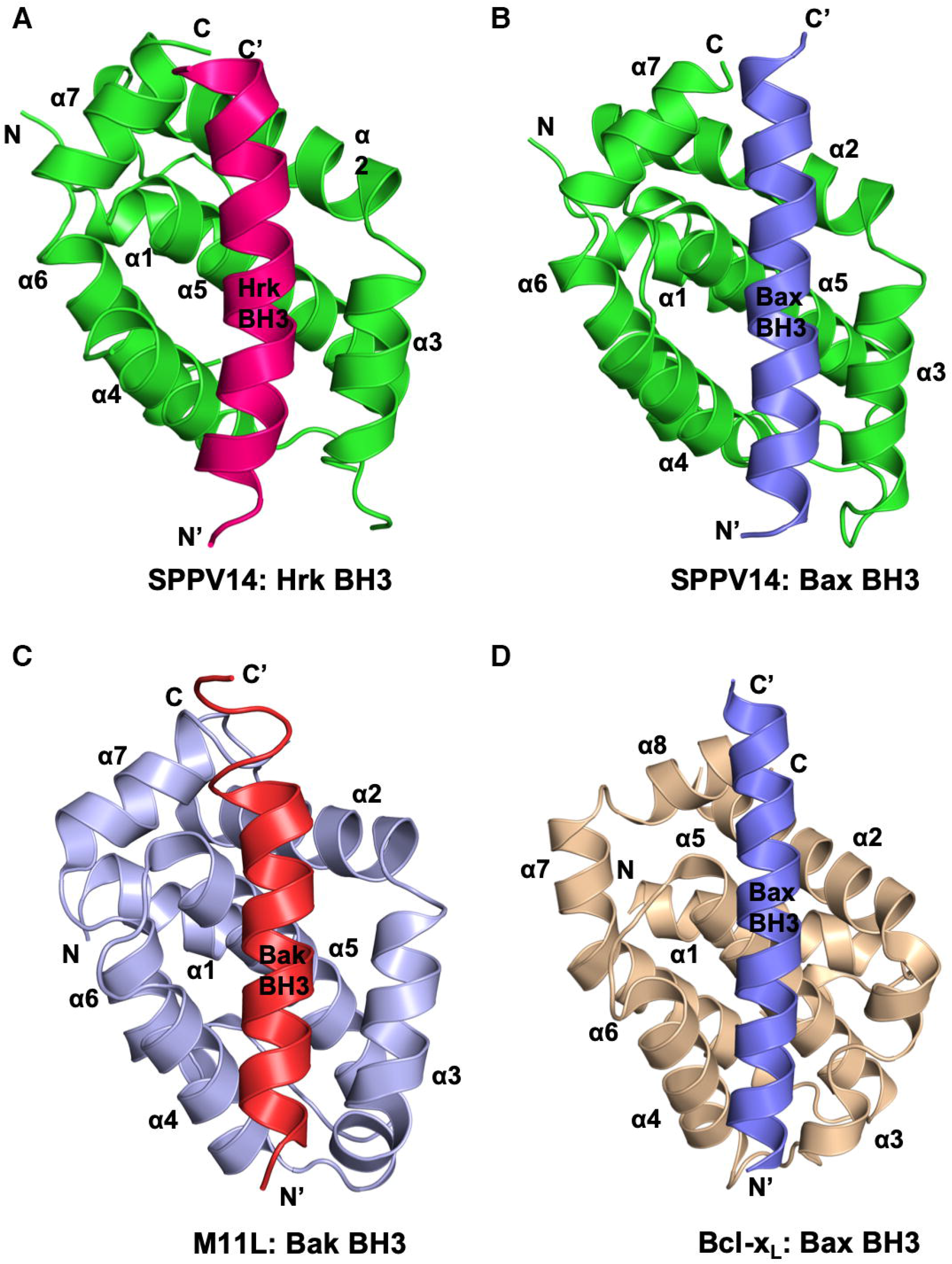
SPPV14 binds to BH3 motif peptides of proapoptotic Bcl-2 proteins using the canonical ligand binding groove. Ribbon representations of Bcl-2 protein BH3 peptide complexes **(A)** SPPV14:Hrk BH3 complex. SPPV14 is shown in green and Hrk BH3 shown in pink and SPPV14 helices labeled α1-α7. The view is into the hydrophobic binding groove formed by helices α2-α5. **(B)** SPPV14:Bax BH3 complex. SPPV14 is shown as in (A) with the Bax BH3 colored slate. The view is as in (A). **(C)** M11L:Bak BH3 complex (PDB ID 2JBY) [18]. M11L is shown in light blue and Bak BH3 is shown in red. **(D)** Bcl-x_L_:Bax BH3 complex (PDB ID 1BXL) [45]. Bcl-x_L_ is shown as sand coloured ribbon and BaxBH3 is shown in slate. The views in C-D are as in (A).

**Table 1:** Interactions of wild-type sheeppoxvirus SPPV14 and two mutants R84A and Y46A with human pro-apoptotic BH3 motif peptides. Measurements were performed using isothermal titration calorimetry. All K_D_ values (in nM) are the means of three independent measurements with standard error. “NB” denotes no binding.

| **Peptide** | **WT**<br>**$K_D$ (nM)** | **R84A**<br>**$K_D$ (nM)** | **Y46A**<br>**$K_D$ (nM)** |
| --- | --- | --- | --- |
| Bak | $48 \pm 1$ | $139 \pm 10$ | $2553 \pm 353$ |
| Bax | $26 \pm 3$ | $182 \pm 15$ | $4645 \pm 393$ |
| Bok | $7580 \pm 643$ | NB | NB |
| Bad | $5197 \pm 533$ | $6700 \pm 806$ | NB |
| Bid | $136 \pm 23$ | $1698 \pm 16$ | NB |
| Bik | $1766 \pm 280$ | $9245 \pm 701$ | NB |
| Bim | $19 \pm 3$ | $44 \pm 3$ | $165 \pm 18$ |
| Bmf | $44 \pm 7$ | $97 \pm 2$ | $151 \pm 18$ |
| Hrk | $39 \pm 5$ | $132 \pm 3$ | $2250 \pm 318$ |
| Noxa | NB | NB | NB |
| Puma | $56 \pm 8$ | $95 \pm 3$ | $988 \pm 142$ |

**Table 2:**
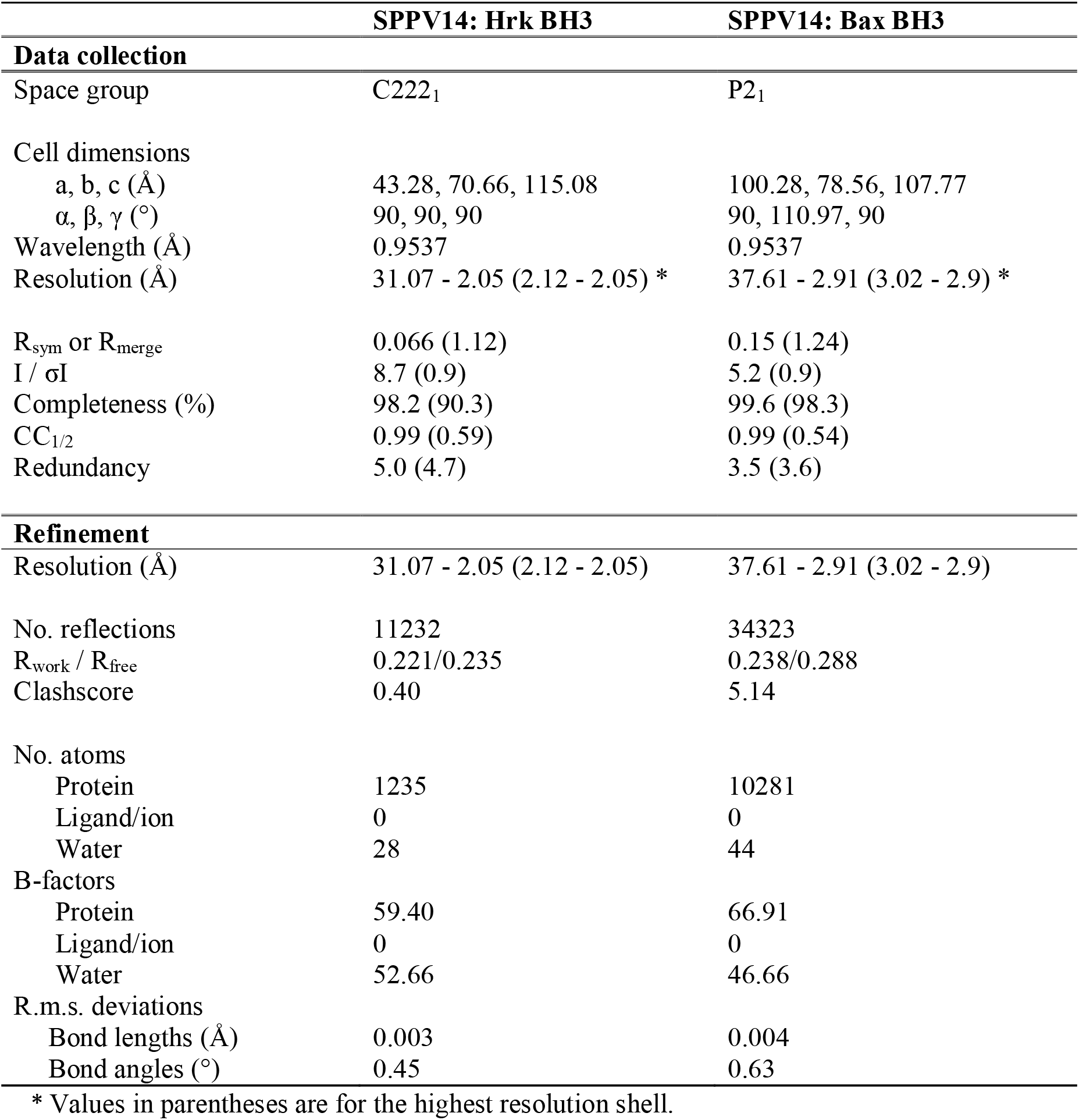
X-ray data collection and refinement statistics.

SPPV14 harbors the canonical ligand binding groove formed by helices α2-α5, which engages Hrk BH3 (Figures 1A, 2A). The mode of binding of BH3 motif peptides observed for SPPV14 is nearly identical to what has been previously observed for M11L:Bak (Figure 2C), with the SPPV14:Hrk complex superimposing onto the M11L:Bak complex with an rmsd of 1.8 Å over the entire SPPV14 chain. The three conserved hydrophobic residues L37, I40 and L44 as well as T35 from Hrk protrude into the SPPV14 binding groove and engage the resident four hydrophobic pockets (Figure 3A). In addition to these hydrophobic interactions, SPPV14 forms an ionic interaction with Hrk via SPPV14 R84 and Hrk D42 carboxyl group. Furthermore, two hydrogen bonds are observed between SPPV14 Y46 hydroxyl and Hrk E43 carboxyl SPPV14 and T78 hydroxyl with Hrk A34 amide.

**Figure 3:**
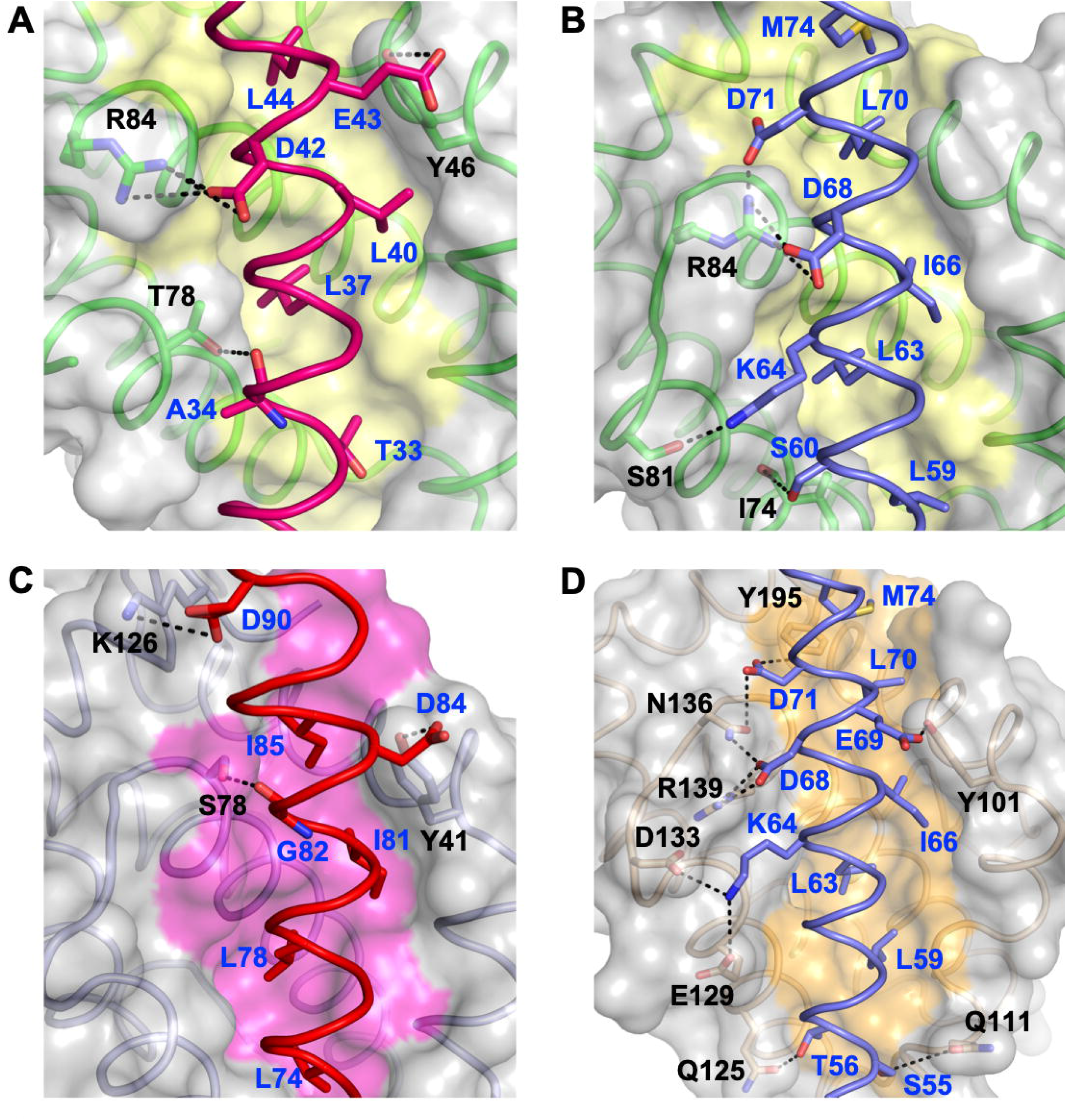
Detailed view of the SPPV14: Hrk BH3, SPPV14: Bax BH3, M11L: Bak BH3 and Bcl-x_L_: Bax BH3 interfaces. **(A)** The SPPV14 backbone, floor of the binding groove and surface are shown in green, yellow and grey respectively, whilst the Hrk BH3 ribbon is shown in hot pink. The three hydrophobic residues of Hrk (L37, L40 and L44) as well as T35 are protruding into the binding groove, and the conserved ionic interaction formed by SPPV14 R84 and Hrk BH3 D42 is labelled, as well as all other residues involved in hydrogen bonds between protein and peptide. **(B)** SPPV14 is shown as in (A) whilst the Bax BH3 ribbon is shown in slate. The five hydrophobic residues of Bax BH3 (L59, L63, I66, L70 and M74) are protruding into the binding groove, and the conserved salt bridge interaction formed by SPPV14 R84 and Bax BH3 D68 is labelled, as well as other residues involved in additional ionic interactions and hydrogen bonds. **(C)** M11L (light blue) is shown as in (C) whilst the Bak BH3 ribbon is shown in red (PDB ID 1JBY). The four hydrophobic residues of Bak BH3 (L74, L78, I81 and I85) are protruding into the binding groove, and the ionic interaction formed by M11L K126 and Bak BH3 D90 is labelled, as well as other residues involved in protein:peptide hydrogen bonds. **(D)** Bcl-x_L_ (wheat) is shown as in (A) whilst the Bax BH3 is shown as slate coloured sticks (PDB ID 1BXL). The five hydrophobic residues of Bax BH3 (L59, L63, I66, L70 and M74) are protruding into the binding groove, and the conserved ionic interaction formed by Bcl-x_L_ R139 and Bax BH3 D68 is labelled, as well as other residues involved in additional ionic interactions and hydrogen bonds. Ionic interactions and hydrogen bonds are shown as dotted black lines and the participating residues are labelled.

The SPPV14:Bax complex was solved by molecular replacement using the SPPV14 model obtained from the SPPV14:Hrk complex and refined to a resolution of 2.9 Å (Figure 1B, Table 2). Clear and continuous density was observed for SPPV14 5-139 and Bax 52-74 (Figure 2B). The four conserved hydrophobic residues L59, L63, I66 and L70 from Bax are nestled in the corresponding four hydrophobic pockets of the SPPV14 binding groove (Figure 3B). Furthermore, Bax M74 also makes contact with the SPPV14 binding groove in a pocket formed by residues C38, V41, I42, F133 and N137. Additional interactions are observed as ionic interactions between SPPV14 R84 and the carboxyl groups of Hrk D68 and D71 as well as hydrogen bonds between SPPV14 S81 hydroxyl and Bax K64 lysyl, and the main chain carbonyl of SPPV14 I74 and Bax S60 hydroxyl group (Figure 3B).

To validate our crystal structures we performed size exclusion chromatography to determine the oligomeric state of SPPV14 in solution. An SPPV14:Bim BH3 complex eluted at a volume commensurate with a heterodimeric species (Figure 4). We next performed structure-guided mutagenesis to modulate the affinity of SPPV14 for BH3 motif ligands (Table 1). Indeed, the SPPV14 mutants Y46A and R84A display attenuated affinities, with SPPV14 Y46A binding BH3 peptides from Bak, Bax, Bim, Bmf, Hrk and Puma with lower affinity and losing binding to those of Bad, Bid, Bik and Bok whilst SPPV14 R84A bound Bax, Bak, Bid, Bik and Hrk BH3 peptides with lower affinities and lost binding to Bok BH3 compared to wildtype protein.

**Figure 4:**
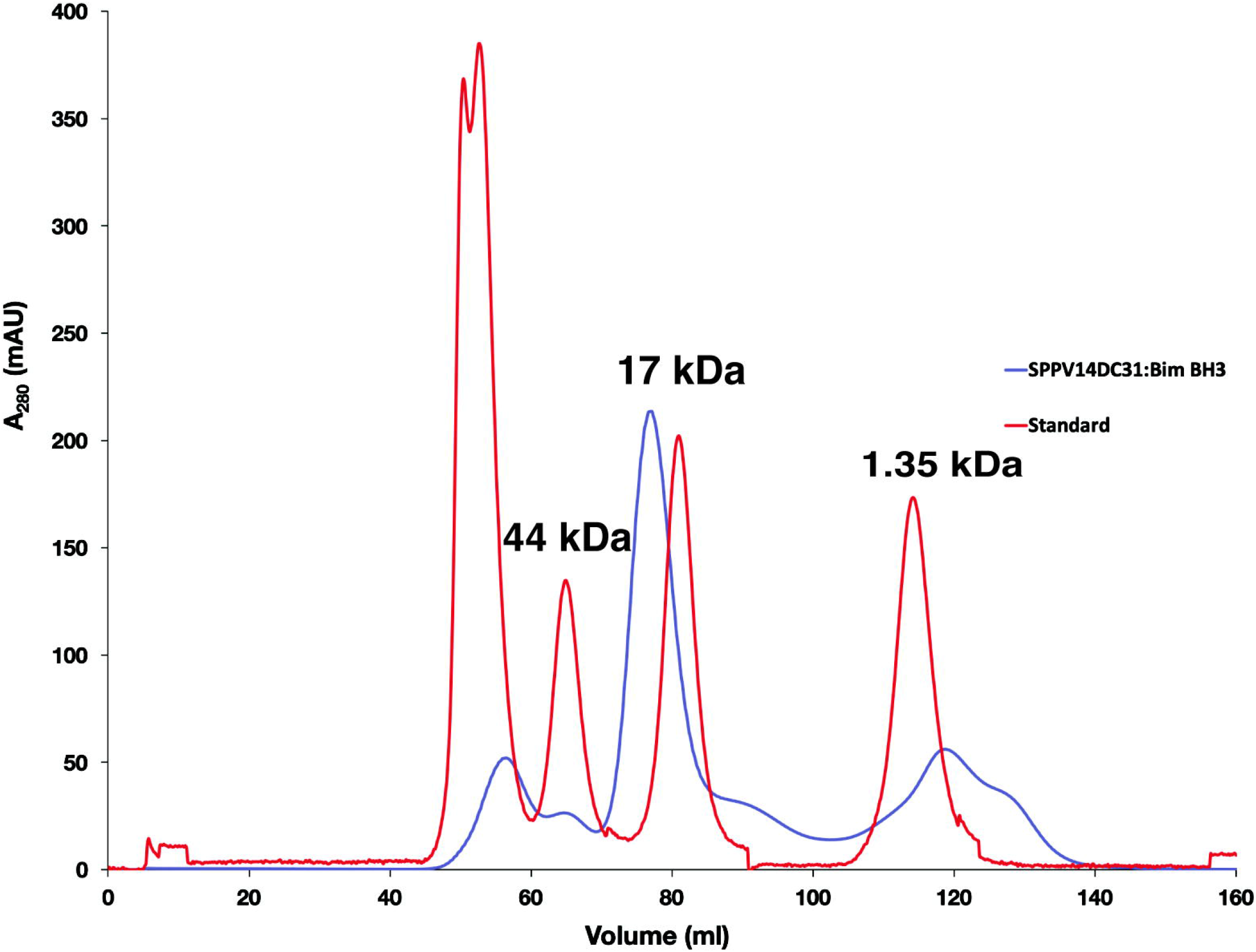
Size exclusion chromatography of SPPV14:Bim BH3 complex. Reconstituted recombinant SPPV14ΔC31:Bim BH3 complex was subjected to size exclusion chromatography using a Superdex 75 16/600 column (blue trace). Molecular weight standards are labelled and indicated on the red trace.

## Discussion

Subversion of premature host cell apoptosis is a frequently used strategy for many large DNA viruses, including the *poxviridae*, to evade the innate immune response [41]. It has previously been shown that sheeppoxvirus harbors the potent apoptosis inhibitor SPPV14, which acts by sequestering BH3-only proteins including Bim, Bid, Bmf, Hrk and Puma as well as Bak and Bax [25]. In addition to the known interactors, we observed previously unreported low micromolar affinities for Bad and Bok BH3 motifs. Previous studies had relied on the use of competition surface plasmon resonance spectroscopy measurements, which could not detect binding affinities weaker than IC_50_ = 2 μM. These findings implicate Bad and Bok inhibition as a potential anti-apoptotic strategy of SPPV14, however considering the comparatively low affinity of the interaction the functional relevance of Bad and Bok inhibition remain to be established.

Having identified potentially new BH3 ligands we now defined the structural basis for BH3 motif interactions with SPPV14 by determining the crystal structures of the SPPV14:Hrk BH3 and SPPV14:Bax BH3 complexes. SPPV14 binds both Hrk and Bax BH3 tightly with K_D_ values of 39 and 26 nM, respectively, and the interactions are mediated by the canonical hydrophobic interactions involving the four hydrophobic pockets in SPPV14 as well as several ionic interactions and hydrogen bonds (Figures 2, 3). A comparison with other virus encoded Bcl-2 proteins reveals that unlike the myxomavirus M11L and vaccinia virus F1L prosurvival Bcl-2 proteins, which both feature low sequence identity to mammalian Bcl-2 [7], SPPV14 forms an ionic interaction using R84 with Hrk D42 or Bax D68 that is reminiscent of the highly conserved ionic interaction between a conserved Arg in the BH1 region of mammalian prosurvival Bcl-2 proteins and the conserved Asp of the BH3 motif (Figure 5A). Such an ionic interaction is seen in all mammalian prosurvival Bcl-2 protein:BH3 peptide complexes exemplified by Bcl-x_L_ R139-Bax D68 (Figure 5A). This Arg:Asp interaction is also observed in the complexes with BH3 peptides of the virus-encoded Bcl-2 homologs A179L [10, 42], BHRF1 [12] and FPV039 [22]. Unlike the canonical ionic interaction [43, 44], the geometric configuration of the ionic bond in SPPV14 is different, allowing SPPV14 to engage BH3 motif in manner that resembles the use of the BH1 motif in mammalian prosurvival Bcl-2 interactions despite the lack of a recognizable BH1 sequence motif (Figure 5A,B). Furthermore, loss of this ionic interaction substantially impacts SPPV14 affinities for BH3 peptide of proapoptotic host Bcl-2 proteins (Table 1), epitomising the behaviour of endogenous mammalian prosurvival Bcl-2 interactions [45].

**Figure 5:**
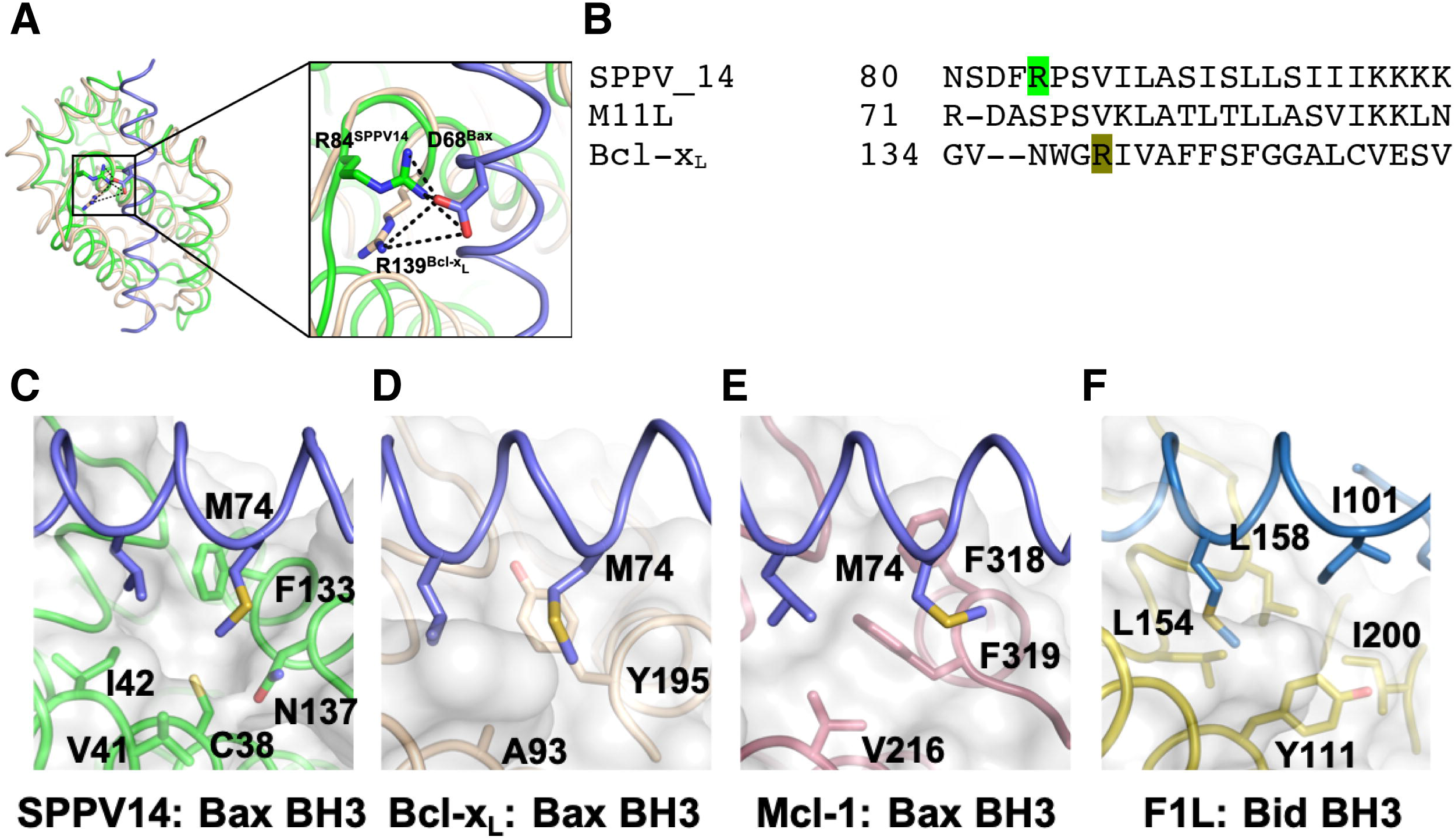
Non-canonical interactions in SPPV14:Bax BH3 complex. **(A)** SPPV14 R84 (green) forms an ionic bond with Bax BH3 D68 (slate) that resembles the canonical ionic interaction between Bcl-x_L_ R139 (sand) and Bax BH3 D68 (slate). For clarity only Bax BH3 from the SPPV14:Bax BH complex is shown. **(B)** Sequence alignment of SPPV14, M11L and Bcl-x_L_ BH1 motifs. Arg residues involved in the ionic interactions with Bax BH3 shown in (A) are coloured. **(C)** Bax M74 protrudes into a fifth pocket in the SPPV14 binding groove formed by C38, V41, I42, F133 and N137. **(D)** Bax M74 bound in the equivalent pocket in Bcl-x_L_ [46]. **(E)** Bax M74 bound in the equivalent pocket in Mcl-1 [46]. **(F)** Bid I101 bound in the equivalent pocket in variola virus F1L [47].

The SPPV14:Bax BH3 complex revealed that an additional residue, M74, engages a fifth pocket in the canonical ligand binding groove besides the four canonical hydrophobic residues that are already accommodated in their respective pockets. This observation mirrors previous findings for Bcl-x_L_ and Mcl-1 [46]. A further example of additional hydrophobic residues from a BH3 motif peptide engaging a pocket is the variola virus F1L:Bid BH3 complex, where I101 from Bid protrudes into a fifth pocket [47]. A comparison of the detailed architecture of these additional pockets (Figure 5C-F) reveals that there is little conservation of residues that form the pockets. In Mcl-1, V216, F318 and F319 are involved, whereas in Bcl-x_L_ it is A93 and Y195. In variola virus F1L, I101 from the Bid BH3 motif engages a fifth pocket formed by Y111, L154, L158 and I200. Interestingly, mutations affecting the binding of Bax M74 to Bcl-x_L_ or Mcl-1 resulted in increased sensitivity to the BH3 mimetic drug ABT-737 [46]. However, the functional consequences of disruption of M74 mediated binding of Bax to SPPV14 remain to be established. We note that binding of the Bax BH3 motif would require substantial structural rearrangements in the context of full length Bax, since in folded cytosolic Bax the BH3 motif is buried and not available for binding to SPPV14 or other prosurvival Bcl-2 proteins [48]. However, the molecular details of such structural changes remain to be defined. In contrast, Hrk is predicted to be an intrinsically unfolded protein, and thus the BH3 motif interactions we observed with SPPV14 are likely to be recapitulated in the context of full length Hrk [49].

Previously several residues were predicted to be located in the binding groove of SPPV14 based on the M11L structure, and thus designated as potential mediators of interactions with BH3 motif ligands [25]. A comparison of the predicted structure with our experimentally determined structures reveal that out of the six predicted residues, all six were indeed found in the SPPV14 binding groove, revealing the high degree of similarity of between the SPPV14 and M11L canonical ligand binding grooves despite an overall sequence identity of 22% [25]. However, whilst six key residues in the binding groove are conserved, the remaining additional BH3 motif contacting residues influence the selectivity or affinity of the viral Bcl-2 for its mammalian targets, as seen by the differential ligand binding profiles and affinities of SPPV14 and M11L. For instance, SPPV14 utilizes Y46 to form a hydrogen bond with Hrk E46, and a similar hydrogen bond is formed by the equivalent tyrosine in M11L, Y41, with Bak D84. Indeed, SPPV14 Y46 is conserved across a number of putative Bcl-2 like proteins. in poxviruses including deerpox, swinepox and rabbitpox [18, 25], as well as in mammalian Bcl-2 proteins such as murine Bcl-x_L_ [43]. In contrast, SPPV14 R84, which forms a salt bridge with Hrk D42 is not recapitulated in M11L. Considering that the salt bridge formed by SPPV14 R84-Hrk D42 mimics other salt bridge interactions seen in many prosurvival Bcl-2 protein:BH3 motif interactions where the prosurvival Bcl-2 protein features broad binding specificity, we speculate that R84 may be a feature that enables the broader binding specificity of SPPV14 when compared to M11L.

SPPV14 is able to directly sequester pro-apoptotic BH3-only proteins as well as Bax and Bak. Whilst numerous large DNA viruses encode for prosurvival Bcl-2 homologs to subvert host cell apoptosis by targeting Bax and Bak [18, 42, 47, 50, 51], mechanisms that do not rely on prosurvival Bcl-2 proteins for Bax/Bak neutralization have also been identified in other viruses. For instance, the herpesvirus human cytomegalovirus encodes for vMIA [52], which has been shown to neutralize Bax by binding in a non-canonical site [53]. Intriguingly, only a short peptide sequence of vMIA is able to mediate the inhibitory effect on Bax. The murine counterpart murine cytomegalovirus (MCMV) encodes the Bax inhibitor m38.5 [54, 55], however the structural basis of this inhibition remains to be established. Inhibition of Bak is achieved by MCMV encoded m41.1 [56, 57]. Strikingly, none of these CMV encoded Bax/Bak inhibitors appear to be multi-motif Bcl-2 homologs, underscoring the importance of Bax/Bak neutralization for the viral life cycle.

In both crystal structures of SPPV14 bound to BH3 motif peptides, SPPV14 appears to be monomeric. Analysis of contacts between SPPV14 chains in the crystal lattices scores all interfaces with a complex significance score (CSS) of 0, suggesting that none of them are biologically relevant. Furthermore, during purification of C-terminally truncated SPPV14 using size-exclusion chromatography, SPPV14 bound to Bim BH3 elutes at a volume corresponding to a heterodimeric species comprising a single chain of SPPV14 (Figure 4). Whilst we cannot exclude the possibility that SPPV14 forms homodimers in a cellular context, our data using recombinantly expressed protein suggest that the active form of SPPV14 is monomeric. Interestingly, a Dali analysis identified DPV022 as the closest structural homolog, even though DPV022 adopts a domain-swapped dimeric topology [20]. Whether or not SPPV14 is able to form similar domain-swapped dimers such as DPV022 or other poxvirus encoded domain swapped dimers such as vaccinia and variola virus F1L is unclear. To date no clear molecular signature that is indicative of an ability to form domain-swapped dimers has been identified in Bcl-2 proteins. It has previously been shown that truncation of the loop connecting helices α1 and 2 in Bcl-x_L_ triggers domain-swapping [58], and in DPV022 and F1L the corresponding α1/2 loops are very short and only 2-3 amino acids in length.

In summary we show that SPPV14 is a monomeric prosurvival Bcl-2 protein that utilizes the canonical ligand binding groove to engage BH3 motif peptides of proapoptotic Bcl-2 proteins. Our structures provide a mechanistic basis to delineate detailed intermolecular interactions and the importance of neutralizing different proapoptotic Bcl-2 proteins for sheeppoxvirus infectivity and proliferation.

## Author contributions

CDS designed and performed experiments, analyzed data and wrote the manuscript. DRB performed experiments, analyzed data and commented on the manuscript. MGH conceived the study, analyzed data and wrote the manuscript. MK conceived the study, designed and performed experiments, analyzed data and wrote the manuscript.

## Acknowledgments

We thank staff at the MX beamlines at the Australian Synchrotron for help with X-ray data collection. We thank the ACRF for their support of the Eiger MX detector at the Australian Synchrotron MX2 beamline and the Comprehensive Proteomics Platform at La Trobe University for core instrument support.

